# Investigating the Intracellular Compartmental Dynamics of miRNAs in Response to Hypoxia

**DOI:** 10.1101/2025.09.18.676626

**Authors:** Vanesa Tomas Bosch, Pierre R. Moreau, Pia Laitinen, Mari-Anna Väänänen, Tiia A. Turunen, Eloi Schmauch, Suvi Linna-Kuosmanen, Minna U Kaikkonen

**Affiliations:** A.I. Virtanen Institute for Molecular Sciences, University of Eastern Finland, Kuopio, Finland; Broad Institute of MIT and Harvard, Cambridge, USA; Computer Science and Artificial Intelligence Laboratory, Massachusetts Institute of Technology, Cambridge, USA

**Keywords:** nucleus, cytoplasm, miRNAs, hypoxia

## Abstract

MicroRNAs (miRNAs) are a class of small non-coding RNAs (ncRNAs) with established functions in cytoplasmic gene regulation, where they interact with mRNA transcripts, impeding their translation—a process known as post-transcriptional gene silencing (PTGS). Recent studies have found that miRNAs and other components involved in PTGS are abundantly present in the nucleus of the cells, where they also play important roles in gene regulation. However, the understanding of their function in the nucleus is not well characterized, particularly in response to cellular stress. We conducted nuclear-cytoplasmic fractionation of primary human endothelial cells (ECs) to assess the distribution of miRNAs during hypoxia. The onset of hypoxia led to changes in miRNA expression in both nuclear and cytoplasmic compartments. Notably, a larger proportion of deregulated miRNAs were identified in the nucleus than in the cytoplasm upon hypoxia exposure. We focused on these nuclear-enriched, differentially expressed miRNAs to investigate their cellular functions and their effects on target mRNAs in different compartments. The data indicated a more pronounced alteration in nuclear mRNA expression compared to cytoplasmic expression, pointing towards a nuclear-specific role for these miRNAs. Several miRNAs influenced target mRNAs associated with pathways of cell proliferation and inflammation. Functional assays under hypoxia showed that miR-5091 and miR-7974 reduced EC cell proliferation, while miR-1246 increased it. Additionally, miR-3662 was found to regulate the secretion levels of specific cytokines, impacting the inflammatory response. Overall, our findings highlight the significant role of miRNAs in the nuclear compartment of ECs in modulating the hypoxic response and provide insights into their multifaceted functions.

## Introduction

MicroRNAs belong to a group of small non-coding RNAs, which primarily exert gene repression by binding to the 3’-untranslated region (3’ UTR) of mRNAs, leading to post-transcriptional gene silencing (Filipowicz et al., 2005). In the canonical biogenesis pathway, miRNA genes are transcribed by RNA polymerase II into a long transcript (Lee et al., 2004), the primary miRNA (pri-miRNA), which is bound and cleaved by the micro-processing complex proteins DGCR8 and Drosha in the nucleus (Gregory et al., 2004), forming a hairpin-structured premature miRNA (pre-miRNA). In the next step of miRNA maturation, pre-miRNA is exported to the cytoplasm through Exportin 5 (Yi et al., 2003) and processed by DICER and TRBP proteins, which cleaves off the hairpin loop resulting in a miRNA duplex (Hutvágner et al., 2001; Wilson et al., 2015). Finally, Argonaute protein (AGO) and other several proteins bind to the miRNA duplex and one strand (known as the guide strand) is kept to form the miRNA-induced silencing complex (miRISC), while the other strand (known as the passenger strand) is degraded (Hu et al., 2009; Meister et al., 2004). Since their discovery in C.elegans, the role of miRNAs in mediating PTGS in the cytoplasm has been extensively studied, where miRNAs act through several pathway. In the initial step, a miRNA recognizes and binds to the miRNA response element (MRE) in the target mRNA. Subsequently, in complex with other proteins, it induces translational repression through mechanisms such as mRNA deadenylation and degradation, 5’-decapping, or ribosome detachment (Huntzinger and Izaurralde, 2011). Surprisingly, although the final steps of miRNA biogenesis occur in the cytoplasm, recent studies have identified mature miRNAs and several of the RISC components, such as Ago2 and TNRC6, to be functional in the nucleus (Gagnon et al., 2014a). In the nucleus, miRNAs can regulate other RNAs -including miRNA biogenesis- through PTGS, as well as mediate transcriptional regulation via epigenetic remodelling at gene promoters or enhancers, or through the interaction with promoter-associated ncRNAs. In addition, nuclear miRNAs have been seen to modulate splicing, and can also serve as an enhancer trigger (El Fatimy et al., 2022; Liu et al., 2018; Roberts, 2014).

Endothelial cells (ECs) play a pivotal role in vascular biology, particularly in processes like angiogenesis and the pathogenesis of atherosclerosis. The response of ECs to hypoxic conditions, a hallmark in both these phenomena, greatly influences cellular behavior and disease outcomes. MiRNAs have been shown to play a key role in hypoxia response (Silina et al., 2023) and have been identified as potential biomarkers or therapeutic targets for vascular disorders. Despite their key recognized role, very little is known of the nuclear and cytoplasmic dynamics and roles of miRNAs. In our previous work, we demonstrated the presence of abundant miRNA species in the nucleus and their significant redistribution between compartments under hypoxic conditions in a mouse endothelial cell line (Turunen et al., 2019). In this study, we aimed to broaden this characterization by conducting a time-course analysis of hypoxia treatment in primary human ECs and provide functional characterization of selected nuclear miRNAs. For the first time, we demonstrate novel functions of nuclear-enriched miRNAs in regulating endothelial cell proliferation and inflammation. These findings suggest a significant role for miRNAs within the cell nucleus and reveal an unparalleled level of complexity in the cellular response to hypoxia.

## Materials and Methods

### Cell culture

Human umbilical vein endothelial cells (HUVECs) were isolated from umbilical cords by collagenase digestion (Davis et al., 2007). The umbilical cords were provided by the maternity ward of the Kuopio University Hospital with the approval from the Research Ethics Committee of the Northern Savo Hospital District. Prior written consent was obtained from the participants and the experiments were performed according to the relevant regulations and Helsinki declaration. HUVECs pool from three different donors were used for obtaining original fractionations upon hypoxia and for miRNA validations. Cells were cultured on fibronectin (10g/ml)-gelatin 0.05% coated plates using endothelial cell growth medium (EGM) supplemented with EGM SingleQuots (CC-3124; Lonza, Basel, Switzerland). Hypoxia was achieved by incubation of the cells in a Ruskinn InVivo2 400 hypoxia incubator with 1% O_2_ and 5% CO_2_.

### Cell fractionation

Cell fractionation was performed according to the protocol by Gagnon *et al*. (Gagnon et al., 2014b) with minor modifications. Briefly, cells were washed with 5mL of warm PBS (#14190169, Fisher), scraped with 3 mL of PBS+0.5%BSA and kept on ice. Next, cells were centrifuged at 1000g for 5 minutes at 4°C. The supernatant was discarded, and pellets were washed with 3mL of PBS+0.5%BSA (# 37525, Thermo Scientific) and subsequently centrifuged at 1000g for 5 minutes. Cell pellet was resuspended in 380 µL of cold HLB, and 5 µL of 100 U SUPERase-IN RNase Inhibitor (#AM2696, Thermo Fisher Scientific) was added prior vortexing. Cells were then incubated on ice for 10 minutes, vortexed and centrifuged 1500g at 4°C for 3minutes followed by 2000g centrifugation for 2 minutes. The supernatant (cytoplasmic fraction) was transferred to a new tube, and 1mL of ice-cold RPS was added. Samples were then vortexed and stored in -20°C for 1 hour, while the pellet (nuclear fraction) was washed with 1 mL of ice-cold Hypotonic Lysis Buffer (HLB) (Gagnon et al., 2014b) three times, by vortexing and centrifugating at 1500g 4°C for 2 minutes followed by 2500g for 2 minutes.

To isolate total nuclear RNA, 1 mL of TRIzol (#AM9738, Fisher) was added to the pellet and stored at -70 °C until RNA purification. Cytoplasmic samples were vortexed after the incubation and centrifuged at 18000g at 4 °C for 15 minutes. Pellet was washed with 1 mL of 75% EtOH, centrifuged 18000g at 4 °C for 5 minutes and air-dried prior addition of 1 mL of TRIzol and stored at -70 °C until RNA purification.

### Transfections

HUVEC cells were seeded in 10 cm plates and transfected at 50% confluency with Mission miRNA mimics (Sigma–Aldrich, St. Louis, MO, USA) listed in Table 1. Mimics were transfected using Oligofectamine (#12252011, Invitrogen, Carlsbad, CA, USA) at a final concentration of 25nM. Supplements were added 4h after transfections, and on the next day cells were washed with PBS and full fresh media was added. Cell fractionation was performed at 48h.

**Table 1.** List of mimic miRNAs used in cell transfections.

| **Product Number** | **Description** |
| --- | --- |
| HMI2235 (Merck) | MISSION MICRORNA HSA-MIR-619-5P |
| HMI2065 (Merck) | MISSION MICRORNA HSA-MIR-5091 |
| HMI2183 (Merck) | MISSION MICRORNA HSA-MIR-5701 |
| HMI0094 (Merck) | MISSION MICRORNA - HSA-MIR-1246 |
| HMI0096 (Merck) | MISSION MICRORNA - HSA-MIR-1248 |
| HMI2145 (Merck) | MISSION MICRORNA HSA-MIR-5585-3P |
| HMI1297 (Merck) | MISSION MICRORNA HSA-MIR-3662 |
| HMI2710 (Merck) | MISSION MICRORNA HSA-MIR-7974 |
| HMI0403 (Merck) | MISSION MICRORNA - HSA-MIR-224 |
| HMI0107 (Merck) | MISSION MICRORNA - HSA-MIR-1257 |
| HMC0002 (Merck) | MISSION(R) MIRNA, NEGATIVE CONTROL 1 |
| HMI0588(Merck) | MISSION MICRORNA - HSA-MIR-454 |

### RNA purification and sequencing

RNA from cell fractionation was purified from TRIzol by adding 10 µL of EDTA 0.5 M (#15575-038, Thermo Fisher Scientific) followed by heating at +65 °C with vortexing until pellet was dissolved.

miRNAs were isolated with RNA Clean & Concentrator kit (#R1018, ZymoResearch), according to manufacturer’s protocol of Purification of Small and Large RNAs into Separate Fractions. On the other hand, purification of mRNA after mimic miRNA transfections was performed with Directzol RNA miniprep kit (#R2052, ZymoResearch), following manufacturer’s instructions.

miRNA-Seq libraries were prepared using NEBNext Small RNA Library Prep Set for Illumina (#E7330L, New England BioLabs), following manufacturer’s instructions, while RNA-Seq libraries were prepared using QuantSeq-Pool Sample-Barcoded 3′ mRNA-Seq Library Prep Kit for Illumina (#44532, Immuno Diagnostics) according to the manufacturer’s protocol. For RNA-Seq, 10 ng of total RNA was used and the libraries were sequenced using paired-end sequencing with Read 1 length set to 70 and Read 2 length set to 22 on an Illumina NextSeq 500 sequencer (Illumina, San Diego, CA).

### cDNA synthesis and qRT-PCR

miRNA reverse transcription was achieved using miRCURY LNA RT Kit (#339340, QIAGEN), according to manufacturer’s instructions. miRNA detection was performed in a LightCycler 96 System using miRCURY LNA SYBR Green PCR Kit (#339345, QIAGEN) and following manufacturer’s protocol. miRCURY LNA miRNA PCR Assay (#339306, QIAGEN) shown in Table 2, which contained forward and reverse PCR amplification primers, were used. Data analysis was performed with Roche LC Software for Cp determination (by the second-derivative method) and melting curve analysis, and *SNORD48* was used for normalization.

**Table 2.** List of miRCURY LNA miRNA PCR Assay used in qRT-PCR.

| **Product Number** | **Description** |
| --- | --- |
| YP00203903 (Qiagen) | *SNORD48(hsa)* miRCURY LNA miRNA PCR Assay |
| YP00205630 (Qiagen) | *hsa-miR-1246* miRCURY LNA miRNA PCR Assay |
| YP00204253 (Qiagen) | *hsa-miR-1248* miRCURY LNA miRNA PCR Assay |
| YP02113278 (Qiagen) | *hsa-miR-1257* miRCURY LNA miRNA PCR Assay |
| YP00204641 (Qiagen) | *hsa-miR-224-5p* miRCURY LNA miRNA PCR Assay |
| YP02118684 (Qiagen) | *hsa-miR-3662* miRCURY LNA miRNA PCR Assay |
| YP00205663 (Qiagen) | *hsa-miR-454-3p* miRCURY LNA miRNA PCR Assay |
| YP02110257 (Qiagen) | *hsa-miR-5091* miRCURY LNA miRNA PCR Assay |
| YP02115478 (Qiagen) | *hsa-miR-5585-3p* miRCURY LNA miRNA PCR Assay |
| YP02106962 (Qiagen) | *hsa-miR-5701* miRCURY LNA miRNA PCR Assay |
| YP02118098 (Qiagen) | *hsa-miR-619-5p* miRCURY LNA miRNA PCR Assay |
| YP02102804 (Qiagen) | *hsa-miR-7974* miRCURY LNA miRNA PCR Assay |

### Proliferation assay

For proliferation assays, 2000 cells/well were seeded in 96-well plates and transfected on the next day as previously described. 48 hours after transfections, cell proliferation assays were performed using CyQuant NF Cell Proliferation kit (#C35007, Thermo Fisher) following manufacturer’s instructions and measurements were done using CLARIOstar Plus microplate reader. After blank subtraction, fold change of transfected mimic compared to mimic control was used as a measure of proliferation.

### Inflammation assay

Cytokine measurements were conducted using the Human Cytokine Array Kit (cat. no. ARY005B; R&D SYSTEMS), according to manufacturer’s instructions. HUVEC cells were transfected with the correspondent mimic miRNA and the cultured media was harvested 24 hours post-transfections for the cytokine array. The immunoblot images were captured and visualized using Bio-Rad ChemiDoc MP imaging system. The intensity of each spot in the captured images was analyzed using the ImageLab 6.1 software (BioRad).

### Data analysis

miRNA-Seq data was analyzed using miRDeep2 (Friedländer et al., 2012) as follows: Initially, adapters were removed with cutadapt (Martin, 2011) and reads were filtered based on length (ranging from 17 to 25 nucleotides) and quality, with reads discarded if less than 90% of their bases had a PHRED score of 30 or higher. Reads containing ambiguous nucleotides (N) were also excluded. Subsequently, the quality of the processed reads was assessed with FastQC (Andrews et al., 2012). Upon confirmation of satisfactory sequence quality, the filtered fastq files were converted into fasta files using the fastq_to_fasta tool (Hannon, 2010). Files mature.fa and hairpin.fa were downloaded from miRbase (Griffiths-Jones, 2006), and hsa_mature.fa and hsa_hairpin.fa files, which contain the mature sequences and hairpins (precursors) for the Homo sapiens species only, were created. The reads were then collapsed using the mapper.pl command in mirdeep2 (Friedländer et al., 2012). Following this step, the reads were mapped against precursor and known mature miRNA species, and their quantity was determined using the quantifier.pl command in mirdeep2. These steps were executed for each sample separately. All sample count files were then aggregated, and duplicate miRNA rows were removed. miRNAs showing a CPM value > 0.5 in at least a third of the libraries were considered expressed. Differential expression was performed on expressed transcripts with HOMER (v4.10) (Heinz et al., 2010) using ‘getDiffExpression.pl’ algorithm with standard parameters.

For RNA-Seq analysis, Fastq files were obtained using bcl2fastq2ConversionSoftware (v2.20, Illumina), and demultiplexed using Idemux (Lexogen). Subsequently, the nf-core RNA-Seq pipeline (version 3.3) with the option “clip_r1: 12”, which instructs Trim Galore to remove 12 bp from the 5’ end of read 1, was used and reads were aligned to GRCh37/hg19 human genome with the STAR aligner (Dobin et al., 2013). Lowly expressed genes have been removed using the filterByExpr function of the EdgeR package (version 3.24.333,34) (Robinson et al., 2010) with a minimum count of 5 reads and minimum total count of 15 reads.

For each microRNA overexpressed, all genes were ranked by their response to miRNA overexpression compared to the miRNA mimic control based on their Wald statistic calculated by the DESeq2 package (version 1.22.235) (Love et al., 2014) using the default parameters. Ranked list were analyzed for gene set enrichment using the fgsea package (version 1.8.038) with the Hallmark gene sets obtained from the Molecular Signatures Database (release 7.139).

### Statistical analysis

Statistical analyses were performed using the GraphPad Prism (v5.03) software. All experiments were performed 2-3 times with three biological replicates per experiment. Data was checked for normal distribution prior performing statistical tests. For data that followed normal distribution, unpaired, two-tailed Student’s t-test was used. Otherwise, nonparametric Mann-Whitney test was used. Statistical significance was evaluated with (*P<0.05, **P<0.005). Results are expressed as mean ±SD.

## Results

### Identification of nuclear and cytoplasmic enriched miRNAs

In order to identify miRNAs enriched in nucleus and cytoplasm, human umbilical vein endothelial cells (HUVECs) were cultured under normoxic or hypoxic (7h and 24h) conditions, followed by cell fractionation and RNA isolation. The small RNA fraction was subjected to microRNA sequencing (miRNA-Seq) and miRNA expression was identified using MirDeep2 (Friedländer et al., 2012). After removing low abundance miRNAs, 654 and 699 miRNAs passed the threshold (CPM>0.5 in at least 3 libraries) in the cytoplasmic and nuclear fractions, respectively (Figure 1A). In addition, we compared our miRNA-Seq data from the fractionation to our previously published HUVEC miRNA-Seq data from whole cell lysates (Linna-Kuosmanen et al., 2020). Although a large fraction (66%) of miRNAs were shared among nucleus, cytoplasm, and whole cell lysate in normoxic conditions, a bigger fraction was shared between nuclear compartment and whole cell lysate, compared to cytoplasmic compartment and whole cell lysate. Interestingly, 6% of miRNAs were only detected in the nuclear compartment, while only 2% were specific to the cytoplasm (Figure 1A). Thus, some of the miRNA enriched in cytoplasmic or nuclear fractions were not detected in the whole cell lysate dataset, emphasizing the advantages of cell fractionation for miRNA detection.

**Figure 1.**
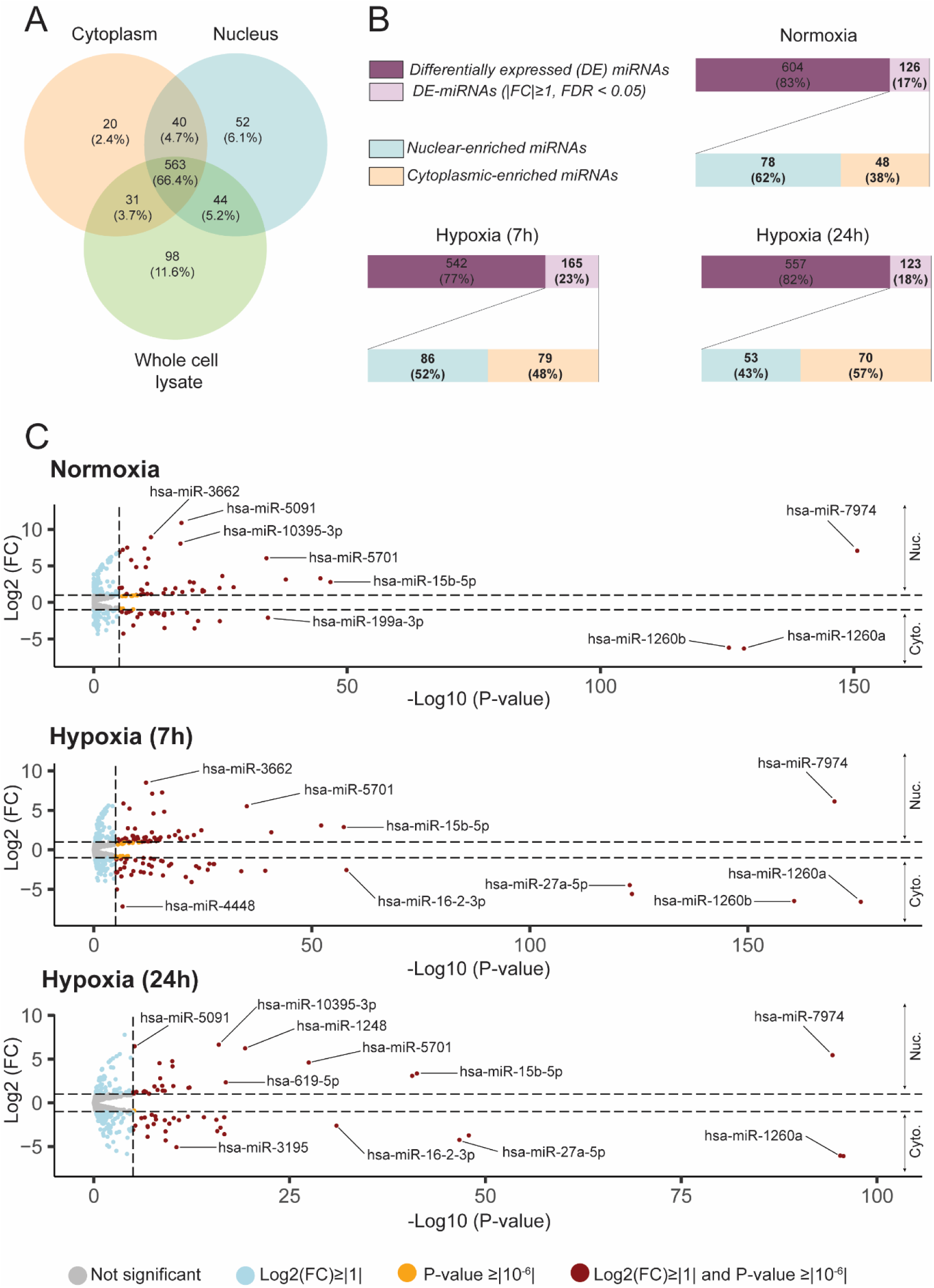
Nuclear and cytoplasmic microRNAs. **A)** Venn diagram comparing identified miRNAs from nucleus, cytoplasm or whole cell lysate of HUVECs cultured in normoxic conditions. **B)** Barplots showing differentially enriched miRNAs and their distribution in nucleus and cytoplasm of HUVEC cells under normoxic and hypoxic (7 and 24h) conditions. **C)** Volcano plots depicting nuclear and cytoplasmic miRNA enrichment in normoxia and hypoxia (7 and 24h). DE, Differentially expressed; FC, Fold-change; FDR, False Discovery Rate; Nuc, nucleus; Cyto, cytoplasm.

Next, we sought to determine nuclear and cytoplasmic enrichment of miRNAs by quantifying the differential expression between nuclear and cytoplasmic samples. We found that, under basal conditions, 730 miRNAs were differentially enriched among nucleus and cytoplasm, from which 126 (17%) were significantly enriched by at least 2-fold in either compartment (Figure 1B). From those, the majority (62%) were enriched in the nucleus, while only 38% of the miRNAs were enriched in the cytoplasm. Upon induction of hypoxia, there was a marked increase in miRNA enrichment within the cytoplasmic fraction. Notably, the range of miRNA enrichment was widespread (Figure 1C). Specifically, miR-1260a and miR- 1260b were among the most enriched miRNAs in the cytoplasm, showing an enrichment of up to 128- fold. Conversely, miR-5091, miR-3662, and miR-10395-3p were some of the miRNAs with the highest enrichment in the nucleus, with levels reaching up to 2048-fold.

### Hypoxia-driven miRNA differential expression in nucleus and cytoplasm

Next, we investigated the effect of hypoxia on the expression of miRNAs in the different compartments. Upon 7h hypoxia, 56 miRNAs were differentially expressed in either compartment (Figure 2A). However, a large majority of them (∼91% - 51/56) were identified in the nuclear compartment and 54% (30/56) were upregulated by hypoxia (Figure 2B). Interestingly, most of these miRNAs were not affected by 7h hypoxia in the cytoplasmic compartment and few of them, such as miR-5091, miR-3648, miR-10396a-3p and miR-10396a-3p, were not detected in the cytoplasmic fraction (Figure 2B, column “C”). In addition, most miRNAs that exhibited changes due to hypoxia in the nuclear compartment did not show similar alterations in the whole cell lysate dataset and several miRNAs were absent from the whole cell lysate dataset altogether (Figure 2B, column “W”). Nuclear and cytoplasmic compartment only shared one miRNA, miR-210-3p, which was upregulated in both compartments and is a well-known regulator of hypoxia (Chan et al., 2012; Turunen et al., 2019). Conversely, only 5 miRNAs were differentially expressed in the cytoplasm and upregulated by hypoxia 7h. From those, only miR-30d-3p and miR-181b- 2-3p showed upregulation in the whole cell lysate dataset as well (Figure 2B, column “W”), while in the nuclear compartment hypoxia did not induce significant changes of these miRNA (Figure 2B, column “N”).

**Figure 2.**
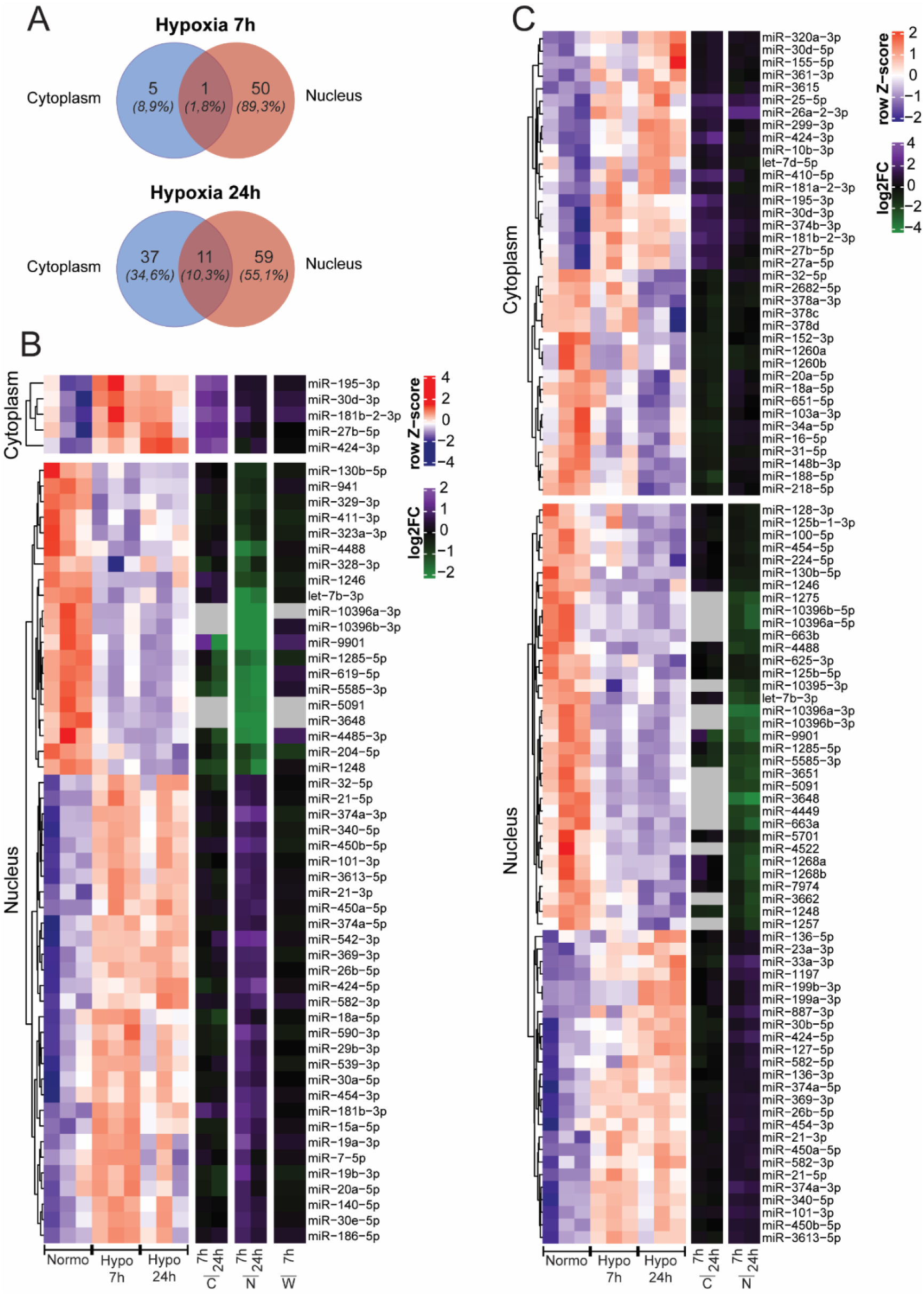
Differential expression of nuclear and cytoplasmic miRNAs upon hypoxic stimulus. **A)** Venn diagram showing the number of differentially expressed miRNAs in nucleus and cytoplasm of HUVEC cells at 7h and 24 h hypoxia. **B)** Heatmap showing differentially expressed miRNAs at 7h hypoxia. Scale bars show mean-centered log2 normalized counts (row Z-score) of the miRNAs found differentially expressed in each compartment, where red and blue indicate higher and lower than mean abundance, respectively. In addition, log2 fold-change of each miRNA in each compartment at 7 and 24 h hypoxia, as well as in the whole cell lysate miRNA-Seq dataset (In Figure 2B) are shown for comparison purposes. In this case, purple and green indicate higher and lower log2FC, respectively. **C)** Heatmap showing differentially expressed miRNAs in the different compartments at 24h hypoxia. Data representation is the same as in Figure 2B. FC, fold-change; Normo, normoxia; Hypo, hypoxia; C, ctyoplasm; N, nucleus; W,Whole cell lysate.

Moreover, we also investigated the effects of 24h hypoxia on miRNA expression (Figures 2A and 2C). Upon hypoxia 24h, 107 miRNAs were differentially expressed, from which 55% were identified in the nuclear compartment. From those, 50% were upregulated and downregulated by hypoxia. As previously seen at 7h hypoxia (Figure 2B), most of the miRNAs identified in the nuclear compartment were not affected by hypoxia in the cytoplasm and 25% (15/60) of them were not detected in the cytoplasmic compartment (Figure 2C, column “C”). Nucleus and cytoplasm shared 11 miRNAs, and interestingly those followed the same trend (upregulated or downregulated) in both compartments (data not shown). Among these, some such as miR-210 (Chan et al., 2012), miR-574-5p (Zhang et al., 2020), miR-542-3p (Wang et al., 2021) and miR-4521 (Xing et al., 2021) have been previously described to be regulated by hypoxia in other cell types. On the other hand, 37 miRNAs were differentially expressed by hypoxia 24h in the cytoplasmic compartment. From those, 50% were upregulated or downregulated by hypoxia and the majority did not show changes in the nuclear compartment (Figure 2C, column “N”), except for few miRNAs which showed upregulation, such as miR-25-5p or miR-26a-2-3p (Kulshreshtha et al., 2007), which has previously been reported to be regulated by hypoxia-inducible factor-1-alpha (HIF-1α).

Furthermore, differential miRNA expression upon hypoxia was broadly distributed and miR-210- 5p and miR-210-3p were highly upregulated at both 7h and 24h in both compartments. The most upregulated miRNAs in the cytoplasm were miR-195-3p and miR-424-3p -both previously described (Chen et al., 2019; Zhang et al., 2022)-, at 7 and 24h of hypoxia respectively, and miR-188-5p was the most downregulated. On the other hand, miR-19a-3p and miR-33a-3p were the most upregulated in the nucleus, at hypoxia 7h and 24h respectively, and miR-3648 was the most downregulated at both time points. Interestingly, although the miRNAs that were differentially expressed in both compartments, followed the same directionality upon hypoxia stimulation, some of them, such as miR-542-3p, had substantial differences of differential expression between the compartments.

Together, these data suggest that miRNAs are differentially regulated by hypoxia in nuclear and cytoplasmic compartments, and, in addition, more miRNAs can be characterized by cell fractionation, in comparison to whole cell lysate sequencing.

### Functional relevance of nuclear-enriched miRNAs upon hypoxia stimulation

In order to explore the cellular effects of nuclear-enriched miRNAs upon hypoxia stimulus, we first selected miRNAs that were enriched in the nuclear compartment (log2FC Nucleus vs. Cytoplasm >0, FDR<0.05) in all time points and differentially expressed (FDR<0.05) by hypoxia 24h in the nucleus. We found 15 miRNAs passing the selected thresholds, from which some have not been extensively characterized (Supplementary Table 1). The expression of these miRNAs in the different compartments and under hypoxia was variable (Figure 3A), with some miRNAs, such as miR-224-5p being highly expressed in both compartments and all conditions, others such as miR-7974 showing a difference of up to 96-fold between the compartments, and in others cases such as with miR-663b and miR-4449, being expressed only in the nuclear compartment and not detected in the cytoplasm. Moreover, the nuclear enrichment upon hypoxia stimulation was also variable, being miR-5091 the most enriched in the nuclear compartment with a ∼256-fold enrichment (Figure 3B), and most of the miRNAs were downregulated by hypoxia at 24h timepoint (Figure 3C), being miR-3651 the most downregulated (∼5-fold downregulation). To better understand the cellular roles of certain miRNAs, we selected a subset based on their nuclear expression levels and previously identified functions. We prioritized miRNAs with higher nuclear expression and those less explored in earlier research. Given that these miRNAs exhibited downregulation under hypoxic conditions, we sought to counteract this effect and overexpress them. To achieve this, we transfected HUVEC cells with mimic miRNAs followed by 24h hypoxia treatment. Subsequently, cells were fractionated and overexpression of the selected miRNAs in each fraction was confirmed by RT-qPCR (Supplementary Figure 1A). Although miRNAs showed the highest overexpression in the cytoplasmic compartment (up to 3x10^5^-fold compared to mimic control), nuclear overexpression was also evident (up to 4x10^4^-fold compared to mimic control). Moreover, nuclear and cytoplasmic markers of a subset of representative samples were checked by RT-qPCR to confirm effective subcellular fractionation (Supplementary Figure 1B).

**Figure 3.**
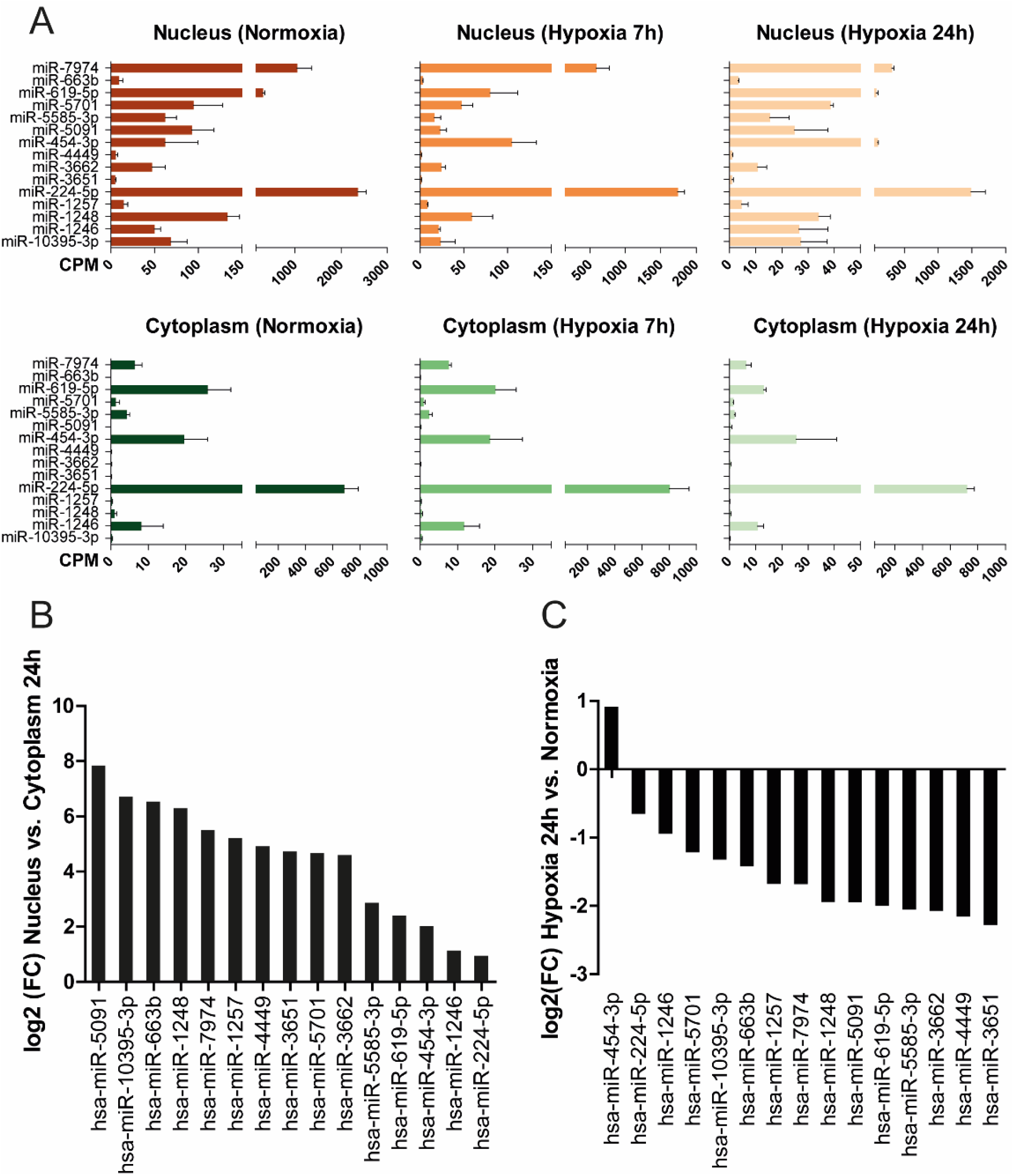
Characterization of nuclear-enriched miRNAs regulated by hypoxia. **A)** Bar plots showing the expression – represented as counts per million reads (CPM)- of the nuclear-enriched miRNAs regulated by hypoxia (24 h). The expression of each miRNA is shown for each compartment in normoxia and hypoxic conditions. **B)** Barplot illustrating the nuclear enrichment of the miRNAs. Nuclear enrichment was determined by differential expression analysis of all nuclear libraries against all cytoplasmic libraries in hypoxic (24h) conditions. **C)** Barplot illustrating differential expression of the miRNAs in the nuclear compartment upon hypoxia 24h. Nuclear and cytoplasmic libraries were considered separately, and differential expression determined between time points. CPM, counts per million reads; FC, fold-change.

Next, to determine the biological pathways of the genes affected by the overexpression of miRNAs, total RNA extracted separately for nucleus and cytoplasm was subjected to RNA-seq, followed by a Gene Set Enrichment Analysis of biological pathways (Figure 4). From our findings, we noted that certain miRNAs, including miR-1246, miR-224-5p, and miR-454-3p, significantly influenced specific cellular pathways in the nuclear compartment without any apparent effect in the cytoplasm, suggesting a distinct nuclear function for these miRNAs. Additionally, for most of the miRNAs studied, some hallmarks showed consistent directional enrichment. For instance, the hallmark “MYC targets”, representing a diverse set of genes regulated by the MYC transcription factor, exhibited negative enrichment upon miRNA overexpression, regardless of the cellular compartment. Interestingly, from all the miRNAs tested, miR-3662 and miR-454-3p showed a positive enrichment in immune-related hallmarks like inflammatory response, IL6-JAK-STAT3 and IL2-STAT signalling pathways.

**Figure 4.**
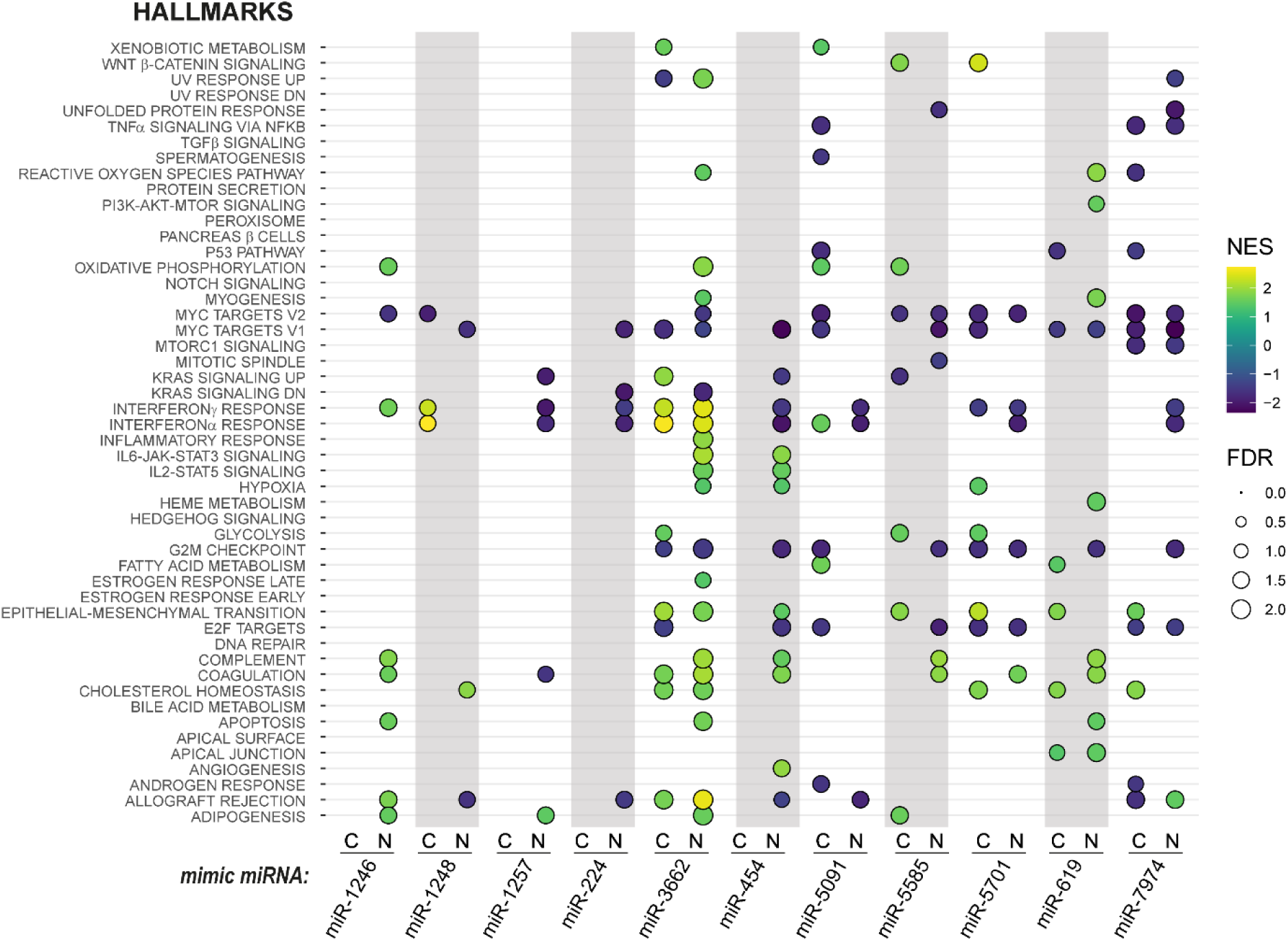
Functional relevance of nuclear-enriched miRNAs upon hypoxia stimulation. Bubble plot showing the pathway enrichment analysis in nucleus (N) and cytoplasm (C) of HUVEC cells transfected with the corresponding mimic miRNA and subjected to hypoxic stimulation (24 h). For clarity purposes, only the most significant bubbles (with an FDR > 1.3) are plotted. NES, normalized enrichment score; FDR, false discovery rate.

### Nuclear-enriched miRNAs are involved in endothelial cell proliferation and inflammation

It is well known that during atherosclerosis progression, hypoxia induces the expression and release of angiogenic factors that stimulate endothelial cell proliferation (Camaré et al., 2017). In addition, dysfunctional endothelium releases various signaling molecules, including adhesion molecules and chemokines, which in turn attract inflammatory cells to the arterial wall (Medina-leyte et al., 2021). Thus, better understanding of these processes is crucial for preventing and treating atherosclerosis and its associated cardiovascular complications.

Since we observed that overexpression of some miRNAs had an effect on several hallmarks that are involved in cell proliferation, such as MYC and E2F transcription factors target genes, KRAS signalling or genes associated with the cell cycle G2M checkpoint among others, we next sought to investigate whether these miRNAs were involved in regulating endothelial cell proliferation. To this end, we performed miRNA overexpression in HUVEC cells subjected to 24h hypoxia, followed by proliferation assay based on measurement of cellular DNA content via fluorescent dye binding (Figure 5A). Interestingly, miR-5091 and mR-7974 showed a significant downregulation in cell proliferation of 40% and 30% compared to mimic control, respectively, in line with the GSEA results showing negative enrichment of gene sets involved in cell proliferation (NES value < 0). On the other hand, miR-1246 significantly increased cell proliferation by 1.8-fold, contrary to the GSEA results, emphasizing the intricate regulatory networks that take place within the cells.

**Figure 5.**
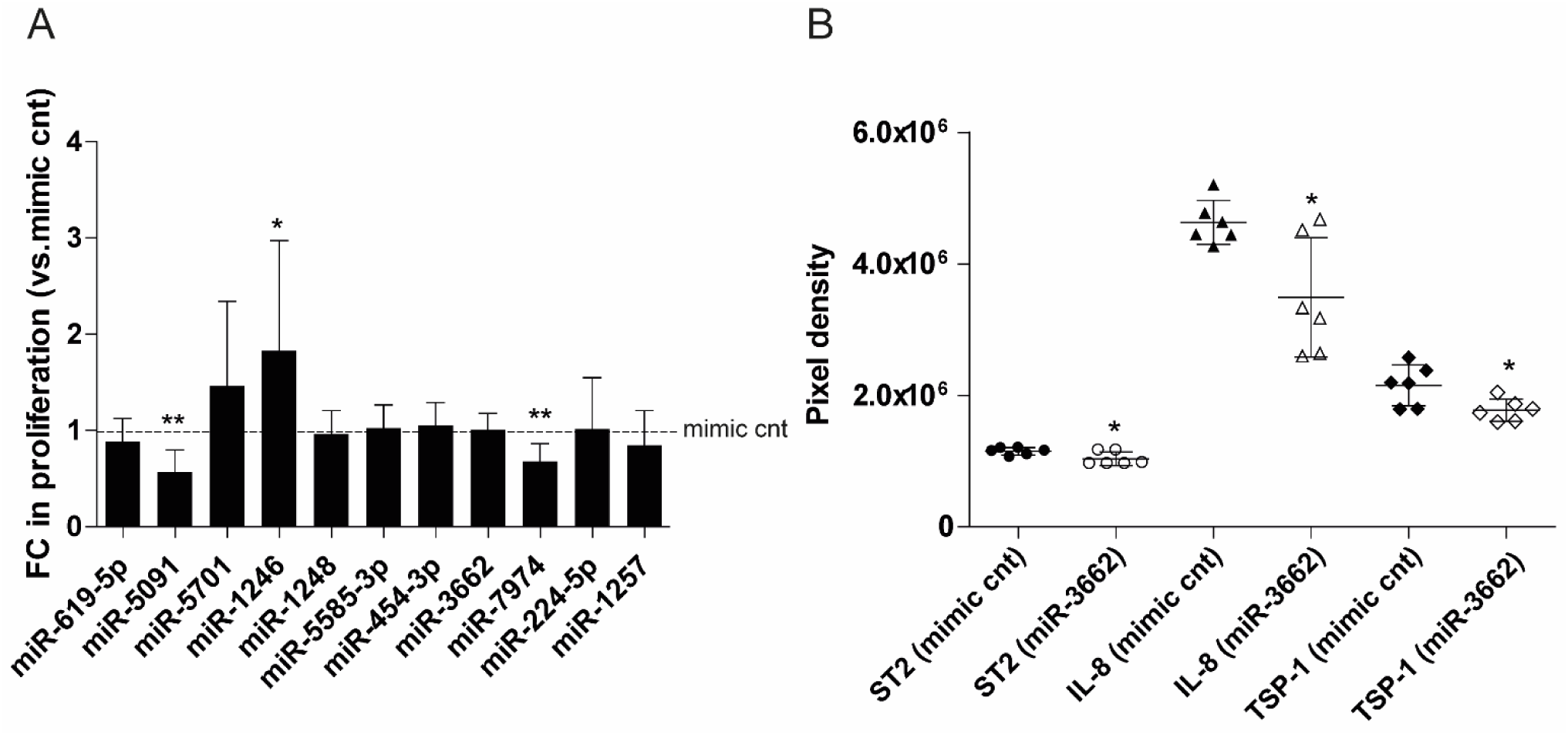
Effect of miRNAs on endothelial cell proliferation and inflammation. **A)** Proliferation assay of endothelial cells transfected with the corresponding mimic miRNAs and subjected to 24h hypoxia. **B)** Cytokine array results showing the pixel density levels detected for ST2, IL-8 and thrombospondin-1 in endothelial cells transfected with either mimic control or mimic miR-3662 and after incubation in hypoxia for 24h. Data was checked for normal distribution before performing statistical tests. Unpaired Student t test (2-tailed) was used for data that followed normal distribution and equal variance. Otherwise, nonparametric Mann-Whitney test was used (mean±SD, n=3, *P≤0.05 and **P≤0.005). FC, fold-change; cnt, control.

In addition, we sought to characterize the regulation exerted by miR-3662 on the modulation of the inflammatory response (Figure 5B). For this, we transfected HUVEC cells with either mimic control or mimic miR-3662, followed by incubation in hypoxia (24h) and cytokine assay. The results showed that from all the cytokines evaluated, miR-3662 caused a modest downregulation in the levels of ST2, IL-8 and TSP-1.

Together, our results show for the first time the role of nuclear-enriched miRNAs in controlling the hypoxic response by affecting cell proliferation and cytokine secretion of human endothelial cells.

## Discussion

In this study, we characterized for the first time the subcellular distribution of miRNAs within human primary endothelial cells in response to hypoxic stimulus. Our miRNA-Seq results revealed an abundant presence of miRNAs in the nucleus and a substantial overlap of miRNAs between the nucleus and cytoplasm, consistent with earlier investigations revealing that approximately 75% of cellular miRNAs were shared across these compartments (Gagnon et al., 2014a; H. Li et al., 2020). Interestingly, we observed a greater concurrence of miRNAs between the nucleus and whole-cell miRNA-Seq data compared to the cytoplasm and miRNA-Seq datasets, and a greater percentage of miRNAs were exclusively detected in the nucleus, highlighting the predominance of miRNAs in this compartment. Hence, based on these findings, mature miRNAs are shuttled back to the nucleus after its biogenesis is completed in the cytoplasm, and are thus generally found to be broadly distributed within both subcellular compartments. However, the specific mechanism of import and the functions of miRNAs in the nucleus are largely unknown.

While earlier investigations have uncovered sequences accountable for directing miRNAs into the nucleus (Hwang et al., 2007), we did not find shared sequence motifs among the nuclear-enriched miRNAs (results not shown). Other studies have proposed that miRNA localization could be contingent upon their interactions with target genes (Roberts, 2014), implying a functional role for nuclear miRNAs in regulating gene expression. In fact, some studies have demonstrated the role of miRNAs in the nucleus of the cells by mediating PTGS of other RNAs (Leucci et al., 2013), as well as by influencing transcription positively or negatively by binding to gene promoters and enhancers, recruiting epigenetic modifying enzymes as well as interacting with promoter associated RNAs (Liu et al., 2018; Stavast and Erkeland, 2019). In addition, other elements accountable for the nuclear localization of miRNAs could be related to their export efficiency to the cytoplasm, the miRNA stability itself or miRNA targeting for degradation. Alternatively, the re-localization of certain miRNAs back to the nucleus could potentially serve as a mechanism to sequester them away from the cytoplasm, allowing for precise control over their impact on translational regulation and mRNA stability. Thus, the distinctive distribution of miRNAs between the nucleus and cytoplasm could serve as a valuable indicator of cellular conditions, such as response to stress, and failing to properly regulate the balance between miRNA localization in the nucleus versus cytoplasm could potentially contribute to cellular dysfunction. In the future, additional research is required to unveil the precise mechanisms and interacting partners responsible for the nuclear import of miRNAs. Hypoxia, characterized by reduced oxygen levels, is a critical physiological condition that cells encounter in various contexts, such as during development, tissue regeneration, and in pathological states like ischemic diseases and cancer. Understanding the molecular responses to hypoxia is of paramount importance, as it can shed light on the intricate regulatory mechanisms that cells employ to adapt to oxygen-deficient environments. One intriguing aspect of cellular adaptation to hypoxia is the differential regulation of miRNAs in the nucleus and cytoplasm, which can have profound implications for gene expression and cellular function. Our findings revealed that hypoxia induces significant alterations in miRNA expression profiles in both the nucleus and cytoplasm, and a modest global downregulation of miRNAs. The observed changes in miRNA expression could be attributed to several factors, such as hypoxia-induced alterations in miRNA export efficiency, modulation of miRNA stability in response to hypoxia or subcellular trafficking mechanisms. Previous studies have reported an overall downregulation of miRNAs in the vascular endothelium during chronic hypoxic conditions, due to a decreased expression and activity of DICER1 enzyme (Ho et al., 2012). Importantly, downregulation of *Dicer1* and consequently, of the processing of pre-miRNAs to mature miRNAs, lead to expression of critical hypoxia-responsive genes regulating the adaptive response to low oxygen availability (Ho et al., 2012). Comparable findings were seen in another study involving cancer cells, wherein the onset of hypoxia led to the suppression of *Drosha* and *Dicer* expression, resulting in a subsequent widespread reduction in miRNA levels and contributing to the progression of the disease (Rupaimoole et al., 2014). Interestingly, a recent study (Pérez-Carrillo et al., 2023) reported a decrease in the amount of mature miRNAs in ischemic cardiomyopathy patients. These findings were the consequence of exportin 5 and DICER downregulation, leading to a decrease in pre-miRNA transport to the cytoplasm and processing to mature miRNAs, respectively.

In our study, sub-cellular fractionation allowed the identification of miRNAs exhibiting distinct expression patterns in each fraction, which were not discernible in whole cell lysate miRNA-Seq data. This underscores the benefits of isolating miRNAs from various cellular compartments when conducting miRNA studies. Moreover, our investigation revealed that hypoxia induced differential expression of miRNAs predominantly within the nuclear compartment, strongly implying a potential functional importance. We selected nuclear-enriched miRNAs differentially expressed by hypoxia for further study of their cellular functions and, importantly, the results showed significant effects on nuclear mRNA expression exerted by these miRNAs, suggesting a possible function of transcriptional regulation in the nucleus of the cells. Particularly, we noticed important effects on cellular pathways related to cell proliferation and inflammation.

Endothelial cell proliferation and inflammation are critical aspects of atherosclerosis in the initiation and perpetuation of the disease (Botts et al., 2021), thus new insights into the molecular mechanisms involved in these processes are critical to identify potential therapeutic targets. Notably, we validated the function of nuclear-enriched miR-1246, miR-7974 and miR-5091 in regulation of cell proliferation, and miR-3662 in regulating inflammation. Prior investigations reported the function of miR- 1246 in the regulation of cancer cell proliferation (Ghafouri-Fard et al., 2022). However, this is the first study to date showing a regulatory role in cell proliferation for miR-7974 and miR-5091, two understudied miRNAs that have received limited research attention and whose functions remained unknown. Additionally, our study uncovered a subtle but significant effect of miR-3662 in several inflammatory pathways, primarily in the nuclear cell compartment. Earlier research has discovered the role that miR- 3662 plays in several cancers, acting as both, pro- and anti-tumorigenic depending on the cancer type (Maharry et al., 2016; Powrózek et al., 2015; Yi et al., 2022; Zhu et al., 2019). In addition, miR-3662 has been shown to interact and decrease the expression of the HIF1α, master regulator of hypoxia (Chen et al., 2018). However, we are the first ones to show its nuclear enrichment in human endothelial cells and its role in regulating secreted levels of ST2, IL-8 and TSP-1. Although our results suggested an upregulation in the inflammatory pathways, miR-3662 overexpression caused a decreased secretion on these cytokines, highlighting the potentially dual role of the identified genes. Indeed, previous studies have described the complex function and association of these cytokines in the development of atherosclerosis. ST2 is a receptor belonging to the interleukin 1 receptor family, which function is to bind to Interleukin-33 (IL-33) and attenuate its functions (Demyanets et al., 2013; Schmitz et al., 2005). Previous research has demonstrated the dual, pro- and anti-inflammatory, role that ST2 exerts depending on the cell type targeted (Altara et al., 2018). Similarly, IL-8 has been described to impact several stages in atherosclerosis development, being both, a pro-inflammatory and anti-inflammatory cytokine (Apostolakis et al., 2009). Endothelial cells are able to synthesize and secrete IL-8, which production is stimulated by other inflammatory molecules and pro-atherogenic flow (Hastings et al., 2007; Yeh et al., 2001). Endothelial cells produce IL-8 in response to inflammatory stimuli, influencing cell proliferation, migration, capillary tube formation, vascular permeability, angiogenesis and adhesion between monocytes and endothelial cells (Gerszten et al., 1999; Li et al., 2003; Luscinskas et al., 1992; Petreaca et al., 2007; Simonini et al., 2000). Concurrently, TSP-1 exhibits a dual role in atherosclerosis as well, depending on the stage of the disease (Stenina and Plow, 2008). TSP-1, involved in cell-matrix interactions and thrombosis, is upregulated in arterial walls following endothelial injury, it facilitates monocyte and macrophage migration (B. Liu et al., 2023; Moura et al., 2007). In addition, TSP-1 also exerts robust anti-angiogenic effects by inducing ECs apoptosis and inhibiting cell migration and proliferation (Jiménez et al., 2000). Our results suggest that hypoxia-mediated downregulation of miRNA- 3662 would cause an increase in ST2, IL-8 and TSP-1. Due to the dual role that these cytokines play in atherogenic context, further studies are needed to investigate the precise mechanism of action and binding partners of miR-3662 in the regulation of these cytokines.

Our prior research has characterized the localization of miRNAs in response to hypoxia in a mouse endothelial cell line and validated the nuclear enrichment of miR-210-3p, the major hypoxia-inducible miRNA, highlighting the relevance of defining miRNA compartmentalization (Turunen et al., 2019). However, it is fundamental to expand this previous research in other cell types and pathophysiological conditions. Hence, our study expands previous knowledge of nuclear miRNAs and paves the way for further research in this field, which ultimately could lead to the development of novel miRNA-based therapies. Studying the role of nuclear miRNAs will open fresh avenues for modulating gene expression, or alternatively could potentially serve as innovative targets in their own regard. While this study provides valuable novel insights, it is not without limitations. The detailed mechanism underlying the effects of nuclear miRNAs is unknown, and additional research is needed to decipher the nuclear targets of these miRNAs. In addition, miRNAs participate in complex regulatory networks: other ncRNAs might also be involved in miRNA-mediated gene regulation, while multiple miRNAs can regulate the same pathways simultaneously, which is not discussed here. Moreover, the interaction of miRNAs with other molecular components of the miRNA biogenesis, such as AGO proteins, as well as transcription factors or epigenetic regulatory enzymes need further investigation. Future studies will need to define what directs the hypoxia-mediated downregulation of some – but not all- miRNAs, as well as their precise mechanism of action. Nuclear miRNAs can act on DNA, RNA or proteins, while cytoplasmic miRNAs regulate mRNA levels. Thus, deeper study will be of great importance to assess their mechanism of action in fine-tuning the hypoxic response, that might even occur via both, canonical and non-canonical mechanisms.

In summary, this study reveals that miRNAs exhibit varying expression patterns in both the nucleus and cytoplasm when subjected to hypoxic stress, providing fresh perspectives on the cellular response to hypoxia and the broader roles miRNAs may play.

## Acknowledgments

We thank the University of Eastern Finland Bioinformatics Center and Biocenter Finland for infrastructure support, and Tuula Salonen for her technical assistance.

## Supplementary Figures

**Figure 1.**
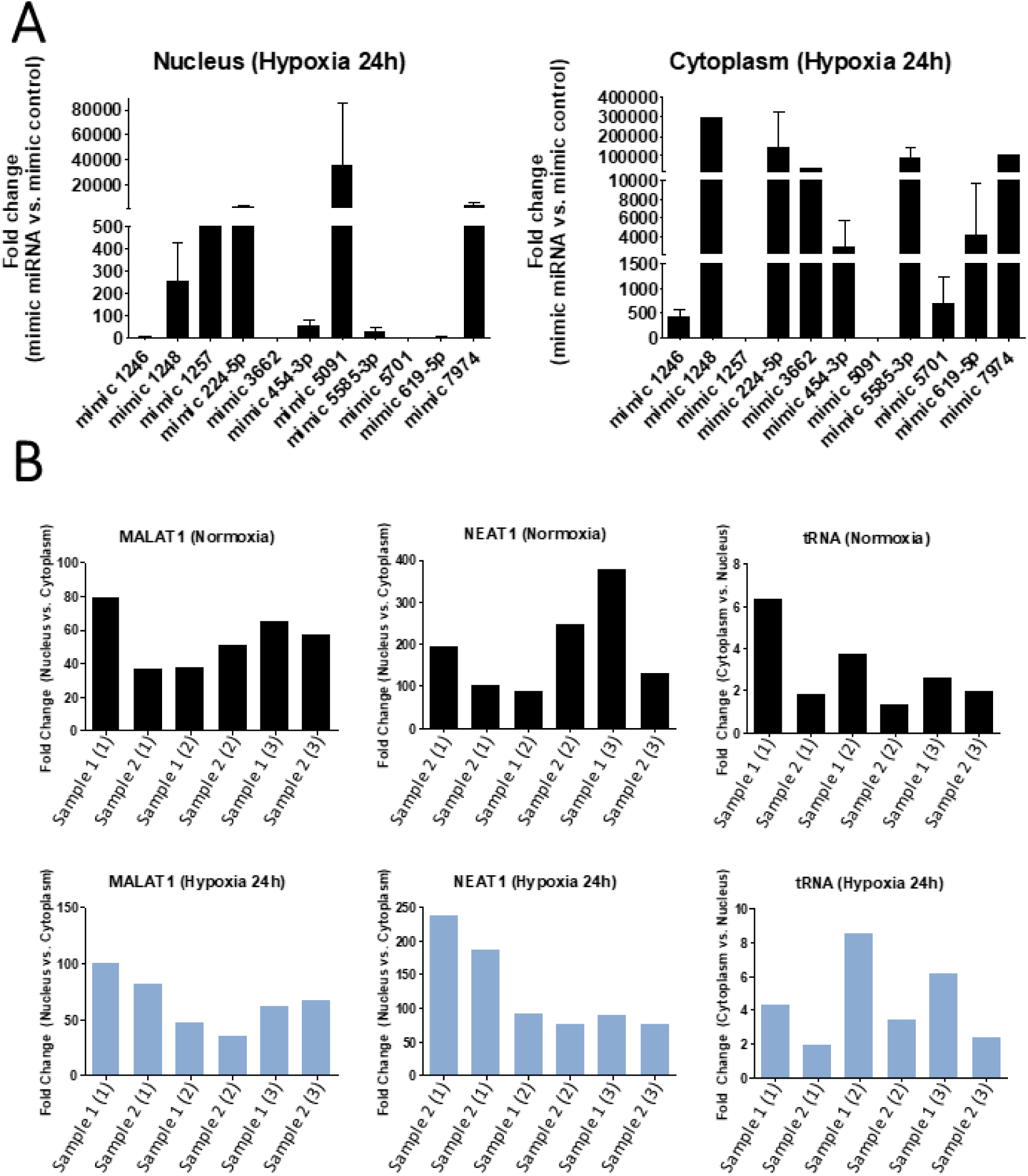
Barplot representing miRNA expression in nucleus and cytoplasm of HUVEC cells transfected with the corresponding mimic miRNAs. Some miRNAs (miR-1248, miR-1257, miR-3662, miR-5091 and miR-7974) were not detected in one or more replicates of the mimic control, hence fold-changes are not plotted. **B.** Barplot representing the enrichment of nuclear and cytoplasmic marker genes after cell fractionation in two representative samples. Numbers in parenthesis represent different replicates.

## Supplementary Table

| miRNA | Function |
| --- | --- |
| hsa-miR-10395-3p | Involved in gastric cancer metastasis and chemoresistance (Luo et al., 2023) |
| hsa-miR-1246 | oncomiR that regulates the expression of multiple cancer-related genes (Ghafouri-Fard et al., 2022; Huynh et al., 2023) |
| hsa-miR-1248 | Exerts a tumor suppressor effect in colorectal cancer (Wang et al., 2022). It is a prognostic biomarker for gliomas (Catanzaro et al., 2022) and it is dysregulated in asthma (Panganiban et al., 2012) |
| hsa-miR-1257 | Role in hepatocellular carcinoma (Gao et al., 2019) and coronary artery disease progression (Xu and Li, 2016) |
| hsa-miR-224-5p | Dysregulated in several types of cancers acting as both, tumoral and tumor suppressor (Córdoba-Lanús et al., 2023; Krasniqi et al., 2021; Li et al., 2022; Y. Liu et al., 2023; Yifei Liu et al., 2022; Naakka et al., 2021; Peng et al., 2021; Rana et al., 2022; Wu et al., 2022; L. Yang et al., 2019; Zang et al., 2020). Dysregulated in amyotrophic lateral sclerosis (Gomes et al., 2023), cystic fibrosis (McKiernan et al., 2021), Behçet's syndrome (Emmi et al., 2022) and patients with reduced coronary flow reserve (James et al., 2022). Involved in osteoporosis (Ding et al., 2023), asthma (Zhuang et al., 2022), hypoxic neuron injury (Liu et al., 2020), adipogenesis (Peng et al., 2013). It has a protective role against hypoxia-mediated cardiomyocyte injury (Mao et al., 2022) acts on pulmonary artery ECs to maintain endothelial function and vascular homeostasis in pulmonary hypertension (Zou et al., 2023). Used as biomarker of breast cancer and endometriosis (Zafari et al., 2021; Zayakin et al., 2023) and involved in chemoresistance in ovarian cancer (Zhao et al., 2014). |
| hsa-miR-3651 | Serves as a predictor for several cancers (Changfeng Li et al., 2020; Pellatt et al., 2016; Ries et al., 2014; Wang et al., 2015; Yerukala Sathipati et al., 2023; Zellinger et al., 2022), plays a role in the development of Alzheimer disease (Wen et al., 2022) and cancer (Yawei Liu et al., 2022) |
| hsa-miR-3662 | Involved in breast cancer (Hosseinpour et al., 2023), hepatocellular carcinoma (Chen et al., 2018), acute myeloid leukemia (Maharry et al., 2016) and lung cancer (Powrózek et al., 2015) |
| hsa-miR-4449 | Involved in schizophrenia, lung cancer (He et al., 2019; Maekawa et al., 2015), multiple myeloma (Shen et al., 2017) and colorectal cancer (Yan et al., 2021) |
| hsa-miR-454-3p | Involved in several cancers (Liao et al., 2021; Hui Liu et al., 2022; Qian et al., 2021; Ren et al., 2019; Shen et al., 2019; Song et al., 2019; S. Wang et al., 2018; X. Wang et al., 2023; Yan et al., 2020), Alzheimer disease (Li and Cai, 2021), angiogenic dysfunction (Cannavicci et al., 2023) and dysregulated in COVID-19 patients (Caixia Li et al., 2020) |
| hsa-miR-5091 | Prognostic biomarker in pancreatic cancer (Liao et al., 2018) and colorectal cancer (G. Yang et al., 2019) |
| hsa-miR-5585-3p | Dysregulated in periodontitis (Baru et al., 2023) patients and in cerebrospinal fluid from patients with intracerebral haemorrhage (Shi et al., 2018) |
| hsa-miR-5701 | Tumor suppressor role in clear cell renal cell carcinoma (Zhao et al., 2021), role in the pathogenesis of Parkinson disease (Prajapati et al., 2018) and its expression correlates with cardiovascular risk factors (Neiburga et al., 2021) |
| hsa-miR-619-5p | Biomarker for lung adenocarcinoma (Feng et al., 2023), esophageal cancer (Okuda et al., 2021) and colorectal carcinoma (Qiu et al., 2016), role in prostatic (Shkurnikov et al., 2017) and gastric cancer (G. Wang et al., 2023), promotes tumor angiogenesis and metastasis (Kim et al., 2020) |
| hsa-miR-663b | Released by cancer cells, promotes angiogenesis in ECs (You et al., 2021a) , involved in several steps of cancer development (Guo et al., 2019; Hong et al., 2020; M. Wang et al., 2018; Yin et al., 2020; You et al., 2021b). Potential biomarker in acute myocardial infarction (Peng et al., 2014), colorectal cancer (Sharma et al., 2021), esophageal adenocarcinoma (Drahos et al., 2016) and epilepsy (Surges et al., 2016) . Role in hypoxia-induced injury in cardiomyocytes (Yu et al., 2020) |
| hsa-miR-7974 | Dysregulated in lung cancer (Dong et al., 2020) |

